# Vascularized human cortical organoids model cortical development in vivo

**DOI:** 10.1101/682104

**Authors:** Yingchao Shi, Le Sun, Jianwei Liu, Suijuan Zhong, Mengdi Wang, Rui Li, Peng Li, Lijie Guo, Ai Fang, Ruiguo Chen, Woo-Ping Ge, Qian Wu, Xiaoqun Wang

**Affiliations:** State Key Laboratory of Brain and Cognitive Science, CAS Center for Excellence in Brain Science and Intelligence Technology (Shanghai), Institute of Biophysics, Chinese Academy of Sciences, Beijing, 100101, China; State Key Laboratory of Cognitive Neuroscience and Learning, Beijing Normal University, Beijing, 100875, China; University of Chinese Academy of Sciences, Beijing 100049, China; Institute for Stem Cell and Regeneration, Chinese Academy of Sciences, Beijing 100101, China; Children’s Research Institute, University of Texas Southwestern Medical Center, Dallas, Texas, USA; IDG/McGovern Institute for Brain Research, Beijing Normal University, Beijing, 100875, China; Beijing Institute for Brain Disorders, Beijing 100069, China

**Keywords:** organoids, cortical development, vascularization, cortical circuit, transplantation

## Abstract

Modelling the neuronal progenitor proliferation and organization processes that produce mature cortical neuron subtypes is essential for the study of human brain development and the search for potential cell therapies. To provide a vascularized and functional model of brain organoids, we demonstrated a new paradigm to generate vascularized organoids that consist of typical human cortical cell types and recapitulate the lamination of the neocortex with a vascular structure formation for over 200 days. In addition, the observation of the sEPSCs (spontaneous Excitatory Postsynaptic Potential) and sIPSCs (spontaneous Inhibitory Postsynaptic Potential) and the bidirectional electrical transmission indicated the presence of chemical and electrical synapses in the vOrganoids. More importantly, the single-cell RNA-seq analysis illustrated that the vOrganoids exhibited microenvironments to promote neurogenesis and neuronal maturation that resembled in vivo processes. The transplantation of the vOrganoids to the mouse S1 cortex showed human-mouse co-constructed functional blood vessels in the grafts that could promote the survival and integration of the transplanted cells to the host. This vOrganoid culture method could not only serve as a model to study human cortical development and to explore brain disease pathology but could also provide potential prospects for new cell therapies for neural system disorders and injury.

## Introduction

In contrast to the rodent lissencephalic cortex, the human neocortex has evolved into a highly folded gyrencephalic cortex with an enormous expansion of the cortical surface and an increase in the cell type and number. The highly developed folded gyrencephalic cortex as well as the alteration of cortical architecture are responsible for the higher cognitive functions that distinguish humans from other species. Animal models, particularly rodents, have provided significant insight into brain development, but the complexity of the human neocortex cannot be fully captured with these models. Therefore, understanding the genetic changes as well as the mechanistic steps that underpin the evolutionary changes that occur during the development of the neocortex in primates may require new model systems.

Organoids have recently been used to study the development and pathological changes of different tissue types, such as the pancreas, liver, kidney, retina etc. (Boj et al., 2015; Huch et al., 2015; Moreira et al., 2018; Morizane et al., 2015; Takebe et al., 2013; Volkner et al., 2016). In addition, several different methods have been developed to mimic neural system development in organoids by differentiating human-induced pluripotent stem cells (hiPSCs) (Jo et al., 2016; Sakaguchi et al., 2015). 3D brain organoids are comprised of multiple cell types that collectively exhibit cortical laminar organization, cellular compartmentalization, and organ-like functions. Therefore, compared to conventional 2D culture, organoids are advantageous in that they can recapitulate embryonic and tissue development in vitro and are better in mirroring the functionality, architecture, and geometric features of tissues in vivo.

In our previous studies, we successfully established an appropriate approach to generate cerebral organoids from hESCs (human embryonic stem cells) or hiPSCs that can recapitulate in vivo human cortical development with the formation of a well-polarized ventricle neuroepithelial structure that consists of vRG, oRG and IPC cells and the production of mature neurons within layers (Li et al., 2017). However, a major limitation in the current culture scheme that is preventing a truly in-vivo-like functionality has been the lack of a microenvironment, such as the vascular circulation. Although they originate from the mesoderm and ectoderm, respectively, the development of vascular systems and neural systems is synchronous in the brain (Paredes et al., 2018). Vascularization is specifically required for oxygen, nutrient and waste exchange and for signal transmission in the brain. Additionally, the blood vessels around the neural stem cells (NSCs) serve as a microenvironment to maintain homeostasis and play essential roles in NSC self-renewal and differentiation during embryonic development (Otsuki and Brand, 2017). The lack of vascular circulation can induce hypoxia during organoid culture and may accelerate necrosis, which consequently interferes with the normal development of the neurons and their potential migration routes (Giandomenico and Lancaster, 2017). To overcome these limitations, some studies have tried to generate vascularized organoids by co-culture of hESCs or hiPSCs with other cell lines (Pham et al., 2018). However, a sophisticated vascular structure that could recapitulate the in vivo vasculature during brain development has not yet been observed in brain-like organoids. Therefore, a more elaborate method is required for establishing better vascularized cerebral organoids to model human neurodevelopment and to look for new therapies for neurologic diseases.

To address this issue, we have developed a three-dimensional culture scheme to generate functional cerebral organoids that are derived from human embryonic stem cells that have been mixed with human endothelial cells in vitro. In our studies, endothelial cells connected to form a well-developed mesh-like or tube-like vasculature in the cerebral organoids (vOrganoids). And the vascularized cerebral organoids mimicked the lamination progress and circuitry formation of human cortical development well and sharing similar molecular properties with human foetal prefrontal cortical cells. Finally, we intracerebrally implanted the vOrganoids into mice and observed that the grafted vOrganoids survived and integrated into the host cortical tissue with relatively normal lamination in vivo. Importantly, the vessels in the vOrganoids connected well with the native blood vessels in rodents to co-build new functional vascularization systems. This vOrganoid culture system serves as a model for studying human cortical development and providing new potential therapeutic strategies to treat brain disorders or injuries.

## Results

### Vascularization in the 3D vOrganoid culture system

Due to the vascularization ability of the human umbilical vein endothelial cells (HUVECs) (Figure S1A), we generated vascularized cerebral organoids (hereon referred to vOrganoids) by co-culturing hESCs with HUVECs. Approximately 3×10^6^ dissociated hESCs and 3×10^5^ HUVECs were plated onto a low cell-adhesion plate, and uniformly sized tight embryoid body-like aggregates formed within the first 7 days. On day 18, the aggregates were transferred to petri dishes for neural induction culture. The resulting 3D aggregates were then re-plated for neural differentiation on day 35 (Figure 1A). The optimized culture conditions allowed these cell aggregates to differentiate and mature up to approximately 200 days (Figures 1B, 1G and 1L).

**Figure 1.**
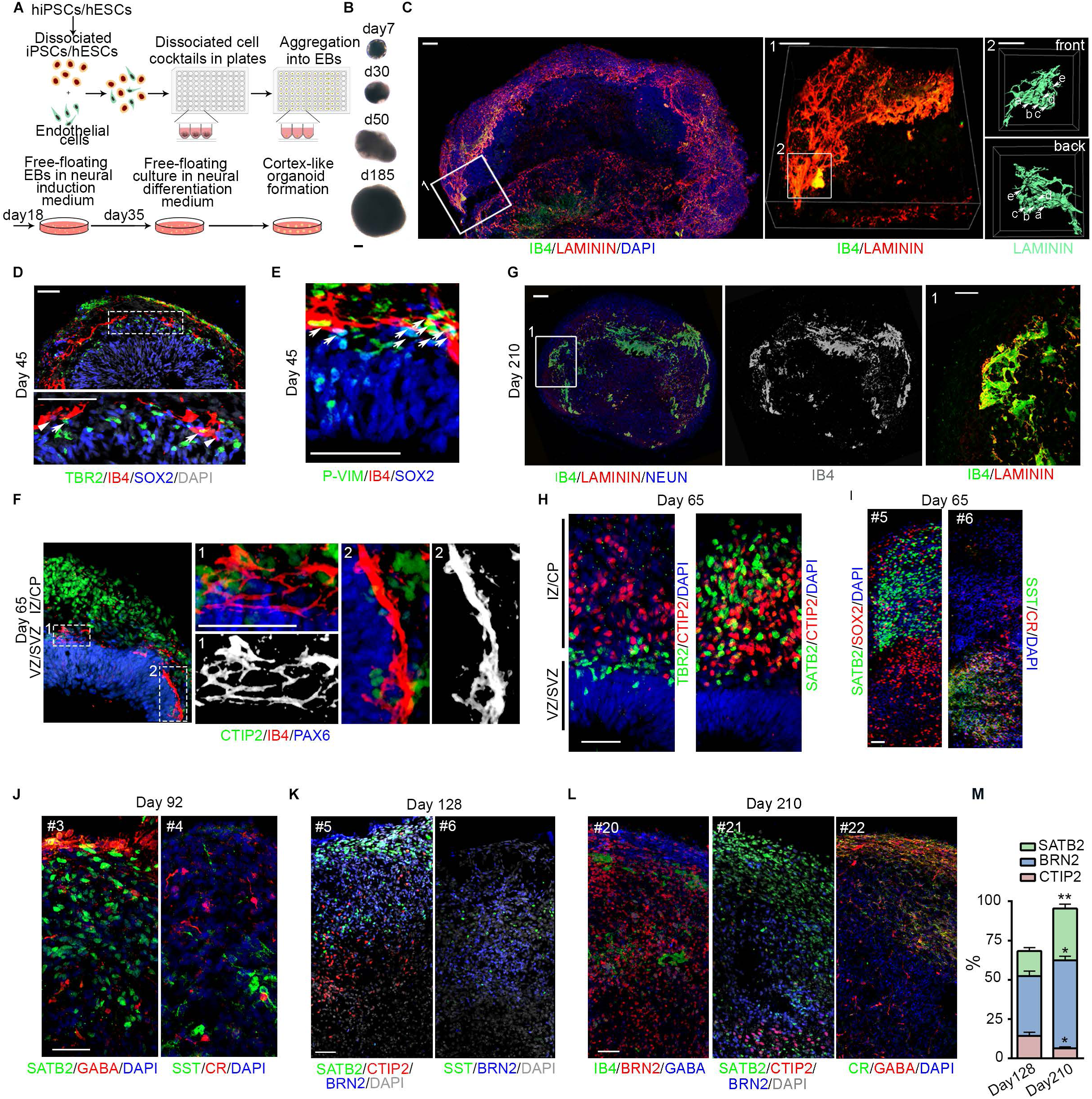
Cerebral vOrganoids with vascular systems recapitulate the cortical spatial lamination. (A) Schematic diagram of the 3D culture methods for generating cerebral organoids with complicate vascular systems. (B) Representative bright field (BF) images of vOrganoids at different stages. Scale bar, 200μm. (C) Whole mount imaging of vOrganoid on day 42. The elaborate mesh-like vascular systems in vOrganoids were displayed by immunofluorescence staining for LAMININ and IB4. Area 1 and 2 outlined in boxes were magnified and 3D reconstructed to depict the complexity of vasculature in vOrganoids. Scale bar, 100μm (left),50um(middle),50um(right). (D) Immunofluorescence staining for TBR2, SOX2 and IB4 to reveal that the vasculogenesis in vOrganoids is coincident with neurogenesis at early stage. TBR2^+^/SOX2^+^/IB4^+^ triple-labeled IPCs were highlighted by arrows, while SOX2^+^/IB4^+^ cells were labeled by arrowheads. Scale bar, 50μm. (E) Immunofluorescence staining for P-VIM, SOX2 and IB4 to demonstrate that IB4^+^vascular systems could also form in the vicinity of P-VIM^+^SOX2^+^oRG cells. Scale bar, 50μm. (F) Immunofluorescence staining for CTIP2/IB4/PAX6 at day 65 demonstrated that the vascular structures progressively extended into newborn neurons as the development of vOrganoids. Scale bar, 50μm. (G) IB4 and LAMININ staining in cerebral vOrganoid at day 210. Scale bars, 100 μm. (H) The spatial lamination of vOrganoids was illustrated by immunofluorescence staining for TBR2/CTIP2(left panel) and SATB2/CTIP2(right panel) at day 65. Scale bar, 50μm. (I) Immunofluorescence staining for superficial layer markers SATB2 and progenitor markers SOX2 to illustrate that cerebral like lamination formed in vOrganoids (left panel); SST and CR staining illustrated that the emergence of interneurons in vOrganoids (right panel). Scale bars: 50 μm. (J-L) Serial cryosections immunostained with pyramidal layer markers and interneuron markers at day 92 (J), day 128 (K) and day 210(L). Scale bars: 50 μm. (M) The percentage of SATB2^+^, BRN2^+^ and CTIP2^+^ cells in cerebral vOrganoids of day 128 and day 210, respectively, n=3 samples. All data are presented as means ± s.e.m. *p < 0.05,**p < 0.01.See also Figure S1.

To visualize the vascularization in the vOrganoids, LAMININ, the major basement membrane glycoprotein of the blood vessel, and Isolectin IB4, a marker for labelling endothelial cells, were both used to reveal the vascular structure. We observed that the endothelial cells were connected into a mesh-like and tube-like structure in 45 day-old vOrganoids (Figures 1C and S1B, Movies S1 and S2). Previous studies have confirmed that the proliferating cells in the VZ/SVZ are, on average, closer to blood vessels (Shen et al., 2008; Tavazoie et al., 2008).

Interestingly, the vascularization in the vOrganoids primarily occurred just above the VZ/SVZ region, where SOX2^+^ RGs and TBR2^+^ IPCs were enriched (Figure 1D), which was in a similar location as the blood vessels in the developing human cerebral cortex of the GW14 (Figure S1C). We also found that the vascular systems were in the vicinity of the mitotic oRGs that were labelled with SOX2 and P-VIM (Figure 1E). All of these results indicated the possible intimate interactions between the vascular system and the VZ/SVZ progenitors during the proliferation. As the vOrganoids developed until day 65, the vascular structures extended into newborn neurons and sustained the growth of the vOrganoids for over 200 days (Figures 1F and 1G). The vOrganoids that were derived from different hESCs lines (H3 and H9 cell lines) showed similar developmental characteristic (Figure S1D), indicating the repeatability of the culture scheme. Compared to the 3D culture without HUVECs, the vascularized organoids were larger with thickened neuroepithelia (Figures S1E-G). Together, in the vOrganoids, the endothelial cells constructed the vasculature in the immediate vicinity with neural progenitors at the early stages and later expanded to the cortical plate-like structures, which may account for the well-developed vascularized organoids with reduced cell death, as a higher degree of vasculature assures sufficient oxygen and nutrient support for cell proliferation and differentiation.

### Recapture subtypes of cells during neurogenesis

To determine whether the vOrganoids could recapitulate cortical spatial lamination, we stained for cortical layer markers on day 65. We found that TBR2^+^ IPCs were adjacent to the early cortical plate (CP), as indicated by the CTIP2^+^ cells (Figure 1H, left panel). Furthermore, later-born superficial layer neurons (SATB2^+^ and BRN2^+^ cells) were more superficially localized to the CTIP2^+^ neurons and accumulated above the PAX6^+^ or SOX2^+^ progenitor cells in the vOrganoids (Figures 1H, right panel; 1I, left panel and S1H).

Since pyramidal neurons and interneurons were both required to form neural circuits, we also observed different subtypes of interneurons when the calretinin (CR) or somatostatin (SST) positive cells were first detectable in the vOrganoids on day 65 (Figure 1I, right panel). Moreover, the layer-specific pyramidal neurons were surrounded by sparse interneurons at the late stages of the 3D culture on days 92, 128 and 210 (Figures 1J-L). The proportion of upper layer neurons increased from 15.94% to 33.04% from day 128 to day 210, and the interneurons were sparsely distributed (Figures 1I-L), indicating that the vOrganoid culture system models the organization of the neocortex in vivo.

Next, we used single-cell RNA sequencing (scRNA-seq) to analyse the cellular and molecular features of the vOrganoids. We collected organoid samples with and without vascular systems at two time points day 65 and day 100, respectively, and analysed a total of 11361 individual cells using 10X Genomics Chromium Single Cell RNA Sequencing. The median number of genes per cell in our study was 3011, and for quality control, the cells with a percentage of mitochondrial genes greater than 0.05 were omitted from the subsequent analysis (Figure S2A). To characterize the cellular heterogeneity of the vOrganoids, we performed a t-distributed stochastic neighbour embedding (t-SNE) analysis by Seurat (Satija et al., 2015) and identified seven major clusters based on the expression of classical gene markers, including RG oRG, IPCs, excitatory neurons, interneurons, microglia and choroid plexuses endothelial cells (Figures 2A-C and S2B). Other than microglia, no glial cells such as astrocytes or oligodendrocytes were clustered in groups (Figure S2C), indicating that this culture system preferably recapitulated the process of neurogenesis in vitro. Some dispersed *RGS5* positive pericytes were detected in our analysis (Figure S2C).

**Figure 2.**
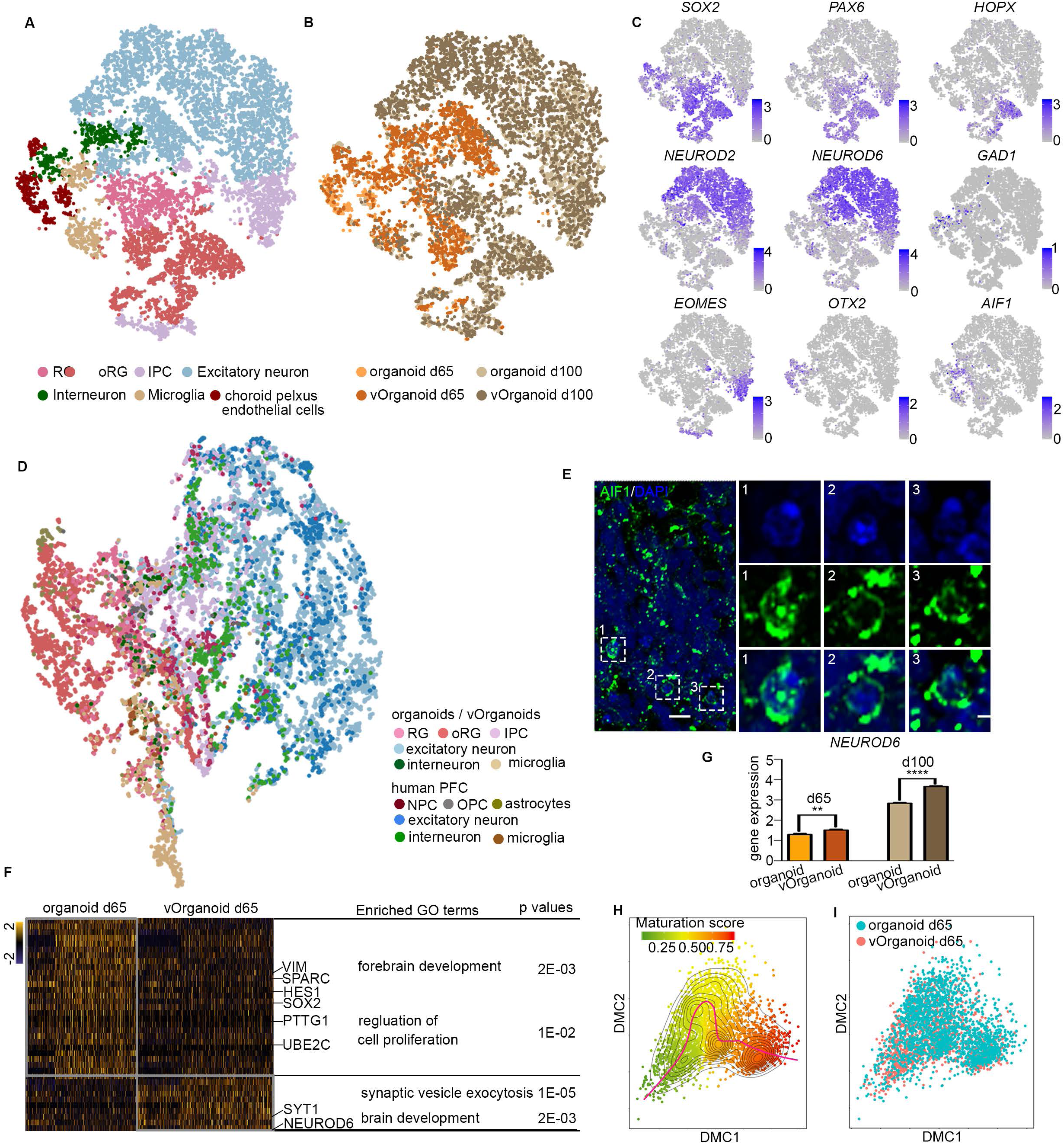
Single-cell RNA sequencing analyses of vOrganoids and human PFC. (A) Visualization of the major cell types in organoids and vOrganoids using t-SNE, colored by cluster. Each dot represents one individual cell. (B) Samples information about the culture time and whether or not with HUVECs was showed using t-SNE, colored by samples information. Each dot represents one individual cell. (C) t-SNE visualizing of the expression of known marker genes. Cells are colored according to gene expression levels (blue, high; grey, low) (D) Organoid cells and embryonic human PFC cells were assigned in tSNE according to the correlations across different cell types. Each cell was colored by cell types. (E) Immunofluorescence staining for AIF1, a specific microglia marker, to illustrate the presence of microglia in organoids. Scale bar, 10μm(left), 5μm(right). (F) Heatmap showing the differential expression genes between organoids with and without HUVECs. Some canonical marker genes were labeled and gene ontology analysis of differentially expressed genes were also listed. (G) The expression levels of NEUROD6, a landmark gene of excitatory neuron, were illustrated in histogram. The NEUROD6 expression levels in vOrganoids are significant higher than organoids. **p < 0.01, ****p < 0.0001. (H) Ordering cells in organoids and vOrganoids along a maturation trajectory. A principal curve was fitted to the dominant diffusion map coordinates. (I) Visualization of the distribution of cells along a maturation trajectory, each dot represents one individual cell. The maturation levels of vOrganoids cells is higher than those in organoids. See also Figure S2.

We then aimed to compare the similarity of the organoids and the human foetal prefrontal cortex (PFC) of GW8-26 (Zhong et al., 2018) at the single-cell level. To identify homologous cell types between organoids and primary tissue, we co-clustered organoid cells and embryonic human PFC cells and assigned individual cells according to the correlations across different cell types, and we found that the same cell types were generally close to each other (Figures 2D and S2D). Microglia, which originate from the mesoderm, are brain parenchymal macrophages (Elkabes et al., 1996). Microglia emerged in our RNA sequencing analysis of the organoids, especially in the vascularized organoids (Figures 2A and 2C). Immunostaining of AIF1, a specific marker of microglia, verified its presence in the vOrganoids, as well (Figure 2E). Notably, we also detected a small cluster of choroid plexus epithelial cells in our data (Figures 2A, 2C and S2C). In vivo, the choroid plexus is mainly responsible for the production of the cerebrospinal fluid (CSF), which serves as a rich source of proteins, lipids, hormones, cholesterol, glucose, microRNA, and many other molecules for the maintenance of CNS functions and plays a role in embryonic neurogenesis (Johansson, 2014). With microglia and choroid plexus epithelial cells, the vOrganoid culture system provides a more complex microenvironment that contributes to the progress of neurogenesis in vivo.

Given the intimate communication between the vascular and neural systems during brain development, we compared the gene expression patterns between the organoids with or without HUVECs to understand how an induced vascular system affects neurogenesis in the organoids. The enriched gene ontology terms showed that the cells in organoid models without a vascular system mainly underwent cellular proliferation, while the synaptic activities were more vibrant in the vascularized organoids (Figure 2F). Additionally, the expression levels of *NEUROD6*, which is a hallmark gene of excitatory neurons, were significantly higher in organoids with a vascular system (Figure 2G), and the maturation analysis (Mayer et al., 2018) illustrated that more cells in the vOrganoids showed high maturation scores (Figures 2H-I and S2E), indicating that the vascular systems in organoids may promote the differentiation and maturation of neurons.

### Modelling functionality maturation during neurogenesis

Several previous studies have illustrated that neurons in cortical organoids without vascular structures are ultimately able to reach mature states as their morphology and functionality progressively mature (Lancaster et al., 2013; Li et al., 2017; Qian et al., 2016). Since vOrganoids partially resemble human cortical development in their molecular and cell subtype aspects, we next investigated the functional characteristics of the neurons with electrophysiological recording. We performed patch-clamp recordings on the slices of the cultured vOrganoids, and Na^+^ currents and spontaneous action potentials were readily recorded in the cells between 90-100 days (Figures 3A and 3B), which is earlier than that of the organoids without HUVECs (Li et al., 2017), indicating that the vascular system in the vOrganoids may accelerate the progression of the functional development of individual neurons in vitro.

**Figure 3.**
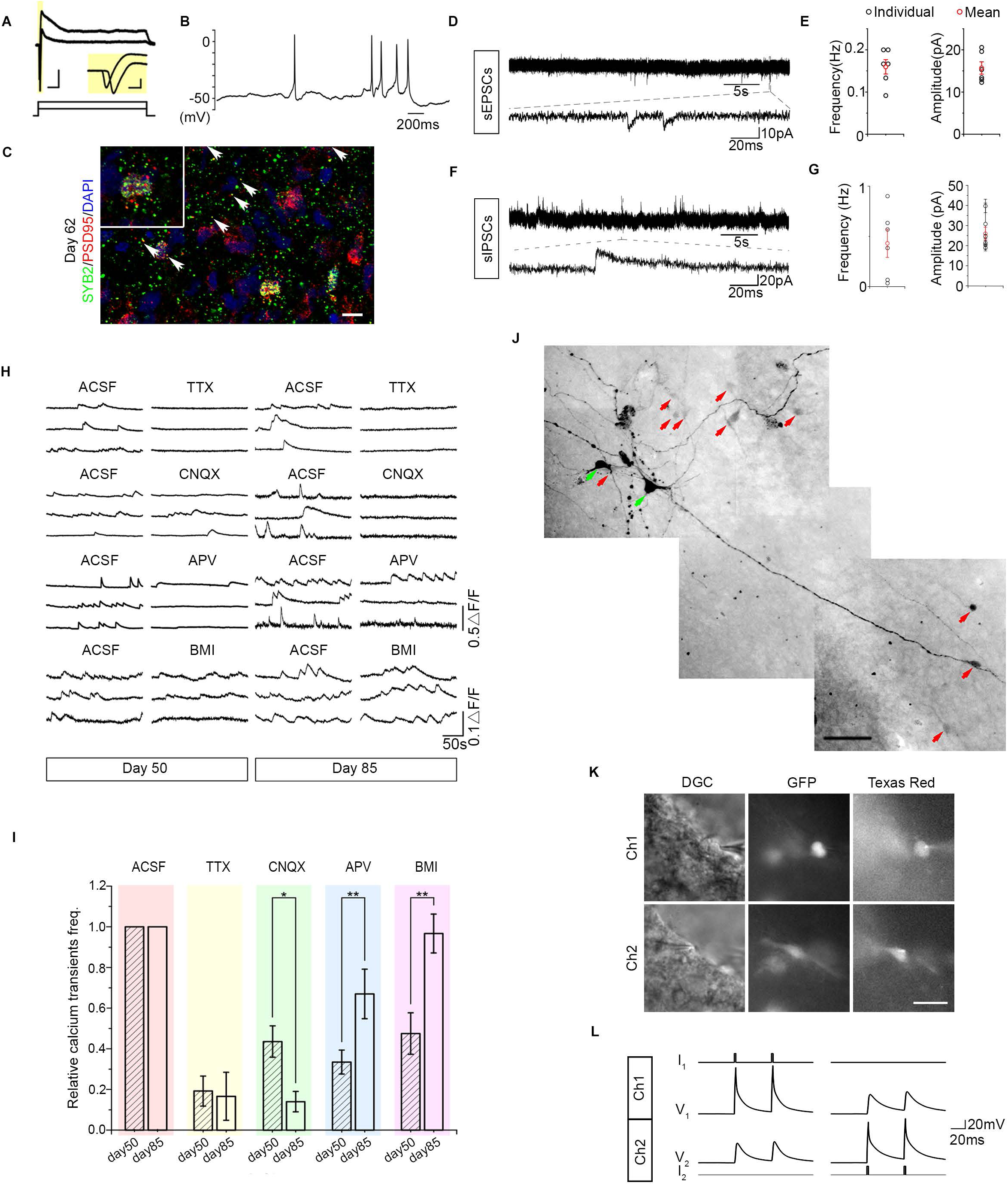
Electrophysiological properties of cells in the vOrganoids at different developmental stages. (A) Representative traces of Na^+^ currents elicited by depolarizing pulses of −20mV or +20mV from a neuron in the Organoids between 90-100 days. Scale bar, 0.5nA, 20ms; 0.5nA, 2ms (insert). (B) An example of spontaneous action potentials recorded from cells of the vOrganoids between 90-100 days. (C) Synapses in vOrganoids revealed by immunofluorescence staining for pre- and post-synaptic markers, SYB2 and PSD95, respectively. Scale bar,10 μm. (D-G) Spontaneous EPSCs (D) and IPSCs (F) were recorded in vOrganoids. The average frequency and amplitude are shown on the right, n=6, 6 cells for sEPSCs (E) and sIPSCs (G). (H) Intracellular spontaneous calcium fluctuations were measured before and after the application of TTX, CNQX, APV and BMI in the vOrganoids at day 50 and day 85, respectively. (I) The effects of different treatments on spontaneous calcium fluctuations were compared between day 65 and day 85 in the statistical results. n=188, 124 cells for day 50 and day 85. All data are presented as means ± s.e.m. * P < 0.05; ** P < 0.01. (J) Coupling pattern was visualized by cell injection. Green arrows indicated the injected cells. Red arrows indicated the cells which were connected to the injected cells. Scale bar, 50 μm. (K-L) Gap junctions between two cells (CMV-GFP labeled) were identified by dual-patch recording. The morphology of cells that dual patched was showed in K. Scale bar, 50 μm in K. The voltage deflections with small amplitudes recorded in one cell while currents were injected into the other cell (L). See also Figure S3.

To build coordinated neural circuits, individual neurons must establish synaptic connections with each other. Therefore, we next focused on the formation of synapses in the vOrganoids. To monitor the formation of the synapses, staining for Synaptobrevin 2 (SYB2) and Postsynaptic density protein-95 (PSD95) were used to identify the presynapses and postsynapses, respectively. We found abundant SYB2 and PSD95 puncta, some of which were colocalized in the vOrganoids on day 62 (Figure 3C). Furthermore, we obtained spontaneous sEPSCs and sIPSCs from the vOrganoids between days 90 and 100 (Figures 3D-G), and the sIPCSc were blocked by the GABAA receptor antagonist bicuculline methiodide (BMI) (Figure S3A). At day 210, dense NeuN^+^/MAP^+^ mature neurons were distributed (Figure S3B), Na^+^ currents and sIPSCs /sEPSCs were captured in these mature neurons (Figures S3C-F), which were indicative of an extensive synaptic formation in the vOrganoids that could sustain a healthy microenvironment for the maintenance of neuronal function.

Intracellular calcium fluctuation imaging is an efficient method for characterizing the functional features of neural activities in cortex-like organoids. Therefore, we performed calcium dye imaging to detect calcium oscillations and observed spontaneous calcium surges in individual cells. When we blocked the action potentials with the application of tetrodotoxin (TTX), a specific blocker of voltage-gated sodium channels, we observed dampened calcium surges in most cases, indicating that spontaneous calcium events in neurons depend upon neuronal activity (Figures 3H and S3G). To further characterize the types of mature neurons, we applied exogenous glutamate and observed more frequent calcium events, indicating that these neurons exhibited glutamatergic receptor activity on day 85 (Figure S3H). Given the predominance of glutamatergic neurons in the aggregates, we combined a pharmacological assay with calcium imaging to further illustrate the different receptor-mediated synaptic transmission functionalities on day 50 and day 85. The neuronal activity in the vOrganoids was reduced during treatments with APV, which is an NMDA receptor antagonist, or CNQX, which is an AMPA receptor antagonist, or BMI, which is a GABAA receptor antagonist, indicating NMDAR-, AMPAR- or GABAAR-mediated synaptic activities appeared after day 50 (Figures 3H and 3I).

In addition to the formation of chemical synapses, we also investigated the electrical synapse (gap junction) in vascularized organoids. To identify the connectivity of neural network, we explored how many cells were connected to one neuron through gap junctions. When we injected neurobiotin into two neurons, 11 coupled neurons were observed (Figures 3J and S3I), suggesting that electrical connections physically exist in the vOrganoids. To directly identify electrical coupling between neurons, we performed dual-patch recording on the vOrganoids. When the dual-patch was established, voltage deflections with small amplitudes could be recorded in one cell while currents were injected into the other cell (Figures 3K and 3L). A bidirectional electrical transmission indicated the existence of gap junctions between these two cells (Figure 3L). Together, these results indicated that the neurons in the vOrganoids could become functionally mature through the emergence of a spontaneous action potential, and functionally matured neurons could later connect to one another based on their abundant chemical and electrical synapse formation and receptor maturation.

### Transplantation of vOrganoids reconstructs the vascular system in mouse cortex

The transplantation of cerebral organoids that are derived from hESCs or iPSCs into injured areas may be a promising therapy for improving neurologic deficits that are caused by trauma or neural degeneration (Daviaud et al., 2018). However, the long-term survival of organoid grafts in a host requires vascularization to satisfy the adequate delivery of oxygen and nutrients. The intracerebral implantation of hiPSC-derived brain organoids into the retrosplenial cortex of immunodeficient mice results in the recruitment of mouse blood vessels that grow into the grafts and support the long-term survival of the cells (Mansour et al., 2018). Therefore, we next tested whether the vascular systems in the vOrganoids could connect to the host brain blood vessels to build a functional circulation system. Sixty-day-old vOrganoids were intracerebrally implanted into a cavity that was made in the S1 cortex of NOD-SCID (nonobese diabetic severe combined immunodeficient) mice (Figure 4A). At 2 months post-implantation (PI), we observed the robust integration and survival of the organoid grafts with a few CASPASE 3 positive apoptotic cells (Figure S4A). To determine whether organoid grafts would exhibit the host brain-like cortical spatial lamination, we conducted immunofluorescent staining of CTIP2 and SATB2, which are deep and superficial layer neuronal markers, respectively. Generally, the SATB2^+^ cells were located above the CTIP2^+^ cells in the organoid grafts, which is consistent with the spatial lamination of the mouse brain. Besides, the implanted grafts displayed slightly folded cortical surface (Figure 4B). We next injected Alexa Fluor 594 dextran into the mouse caudal veins to observe the blood flow in the organoid grafts by live two-photon microscopy. Interestingly, a steady blood flow was captured in the organoid grafts, indicating the formation of a functional vascular system between the grafts and host (movies S3). The three-dimensional reconstructed images also show the morphology of the blood vessels in the grafts (Figure 4C, movie S4). In addition, to further demonstrate that endothelial cells in the grafts had reliably mixed into the new vascular systems, Dextran and LAMININ were immunofluorescently stained to label blood vessels, while HUN was used to distinguish the vOrganoid grafts from the host rodent cells. We observed that human and mouse endothelial cells co-existed in the blood vessels in the grafts 2 months after the implantation (Figures 4D and S4B).

**Figure 4.**
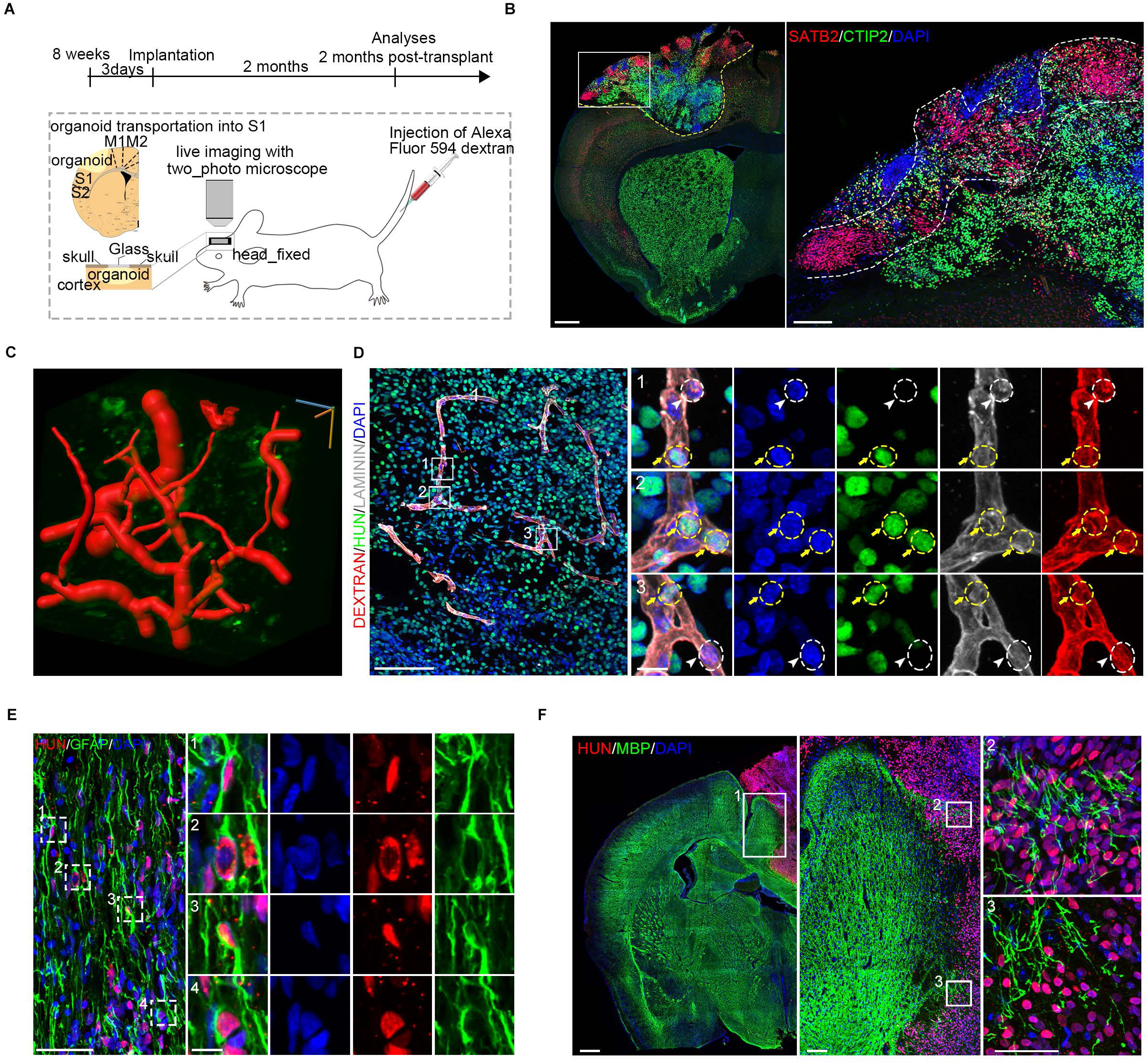
The vOrganoids play important roles in the reconstruction of vascular systems after transplantation. (A) Schematic diagram to demonstrate the vOrganoids implantation protocols in our studies. The vOrganoids were transplanted into the S1 cortex of NOD-SCID mice. (B) Immunofluorescence staining for classical cortical layer markers, CTIP2 and SATB2, at 2 months post implantation. The SATB2 positive cells were roughly located above CTIP2 positive cells, which was consistent with human cortical lamination. The boxed area was magnified at right panel. Scale bar, 500 μm (left), 50 μm (right). (C) 3D reconstruction of blood vessels located at vOganoids grafts. The red tubular structure was blood vessels. Scale bar was labeled at three dimensions. Scale bar, 30μm. (D) Immunofluorescence staining for LAMININ, Dextran and HUN were performed to confirm that human endothelial cells derived from vOrganoids (labeled by yellow circles and yellow arrows) and mice endothelial cells derived from hosts (labeled by white circles and white arrowheads) both played roles in reconstruction of functional vascular systems. Scale bar, 100 μm (left), 10 μm (right). (E) Immunofluorescence staining for GFAP and HUN to illustrate that whether vOrganoids derived astrocytes present in grafts. A few of HUN positive astrocytes was labeled by boxes and magnified at right panels. Scale bar, 50 μm (left), 10 μm (right). (F) Immunofluorescence staining for MBP and HUN to illustrate that whether vOrganoids native myelinization occur at the graft regions. The boxed areas were magnified. Scale bar, 500μm (left), 100μm (middle), 50μm (right).See also Figure S4.

Considering the supporting roles of glial cells in normal neural activities and the integration of grafts into host tissue, we next examined the level of gliogenesis within the organoid grafts. Abundant GFAP^+^ astrocytes were present in the graft regions, and some of them were HUN positive, which suggested that some vOrganoid native astrocytes and host astrocytes played a role in supporting or regulating neural activities (Figure 4E). Furthermore, we also detected signs of myelination in the grafts with staining for myelin basic protein (MBP). However, consistent with the observation that very few oligodendrocytes were found in the cultured vOrganoids with scRNA-seq, only sparse MPB positive signals were observed at the implantation junction of the grafts, but no myelination was detected in the core regions (Figure 4F), which is consistent with previous studies (Mansour et al., 2018). These results suggested that the myelinated fibres in the organoid grafts were primarily derived from the host brain and merely intruded into regions of the implantation junction and that barely any came from the organoids at the early implantation stages.

## Discussion

Neurovasculature, which is the circulatory system in the brain, is a key component of the NSC microenvironment, which provides oxygen and nutrients supports to the brain while removing waste metabolites. The proper formation and function of the central nervous system (CNS) relies on mutual crosstalk between the nervous and the vascular systems (Otsuki and Brand, 2017; Paredes et al., 2018; Tan et al., 2016). Brain organoids are promising models for investigating the development of the human brain in addition to mental disorders. However, in current culture systems, oxygen and nutrients are hampered in reaching the centre of the organoids due to impairments in circulatory system; therefore, cellular necrosis occurs in the centre. Cellular necrosis in organoids, in turn, heavily limits the continuous growth and long-term maintenance of functional cells in organoids. Here, we have established a highly reproducible system for the generation of vascular cerebral vOrganoids by co-culturing hESCs with HUVECs with minimal introduction of extrinsic signals. In this system, endothelial cells connected to form a mesh-like and tube-like vascular structure in the vOrganoids. In the early culture stage, the vascular structures preferred to be adjacent to neural progenitors and were located precisely above the VZ/SVZ region, which is consistent with the location of these vessels in vivo. Endothelial cells have been proven to secrete soluble factors such as vascular endothelial growth factors (VEGFs), angiopoietin-1 (Ang1) and angiopoietin-2 (Ang2) that promote the self-renewal and differentiation of the V-SVZ progenitors (Androutsellis-Theotokis et al., 2009; Jin et al., 2002; Sun et al., 2006). Additionally, the VZ/SVZ is more densely vascularized than its neighbouring brain regions during development (Culver et al., 2013; Kazanis et al., 2010). Several studies also indicate that neural cells in the neighbouring locations could participate in feedback mechanisms to regulate the growth and properties of the cells in the vasculature (Paredes et al., 2018). Therefore, our vOrganoid systems, to a certain extent, recapitulate the in vivo interaction between vascular cells and neural stem cells/progenitors during neurogenesis. In the vOrganoids, the endothelial cells may secrete essential extrinsic signals that facilitate neurogenesis. However, although no real blood cells are present in the vascular structure, hollow tubing structures could possibly supply even greater levels of oxygen and nutrients for neural progenitors and neurons, which may promote cell proliferation and differentiation and prevent cell death. These two aspects may account for the larger size and earlier maturation of neurons in the vOrganoids compared to the organoids in which HUVECs were not added. Given the limitations of traditional organoid culture, our 3D cerebral culture with vascular systems satisfactorily solved the problems of insufficient oxygen and nutrients supports in the organoids, thus permitting the better growth and functional development of the vOrganoids. Thus, our culture systems may be an effective route to improve the traditional methods of neural differentiation.

The vOrganoids mimic cortical development in vivo and contain RG, oRG, excitatory neurons, interneurons and microglia as their major cell types, which are assigned to the appropriate localizations. In contrast to the cell composition of the human prefrontal cortex, sporadic astrocytes, pericytes, and clustered choroid plexus epithelial cells were detected by scRNA-seq in the vOrganoids. Unfortunately, we did not find clustered endothelial cell in the vOraganoids, but this may be explained by the mild dissociation method we used that could barely break the tight junctions between endothelial cells. In vivo, the choroid plexuses are heavily vascularized tissues that are suspended in the ventricles and responsible for secreting the CSF (Lehtinen et al., 2013). RG can intercalate or penetrate the ependymal cell layer and extend a primary cilium into the ventricle to communicate with the ependymal cell/CSF niche (Conover et al., 2000; Mirzadeh et al., 2008; Shen et al., 2008). The CSF represents a single-access route for blood-borne or vascular endothelial cell signals to reach the V-SVZ NSCs. Due to the presence of endothelial cells, pericytes and choroid plexus epithelial cells, the microenvironments for neurogenesis in the vOrgnoids may be more similar to those of the in vivo state compared to other models.

Pyramidal excitatory neurons and interneurons in the vOrganoids were aligned in layers that were similar to those of human neocortical lamination. With this elaborative cytoarchitecture, the neurons gradually matured with synaptic connection formation. We also found a transition from an NMDA receptor to an AMPA receptor mediated transmission of excitatory synapses in our culture system, indicating a maturation process of the neurons that contributes to the formation of a functional circuit (Yu et al., 2014). Together, our vOrganoids that were derived from hESCs with endothelial cells mimic human cortical developmental not only in their cell types and cellular lamination but also in their circuit formation.

A variety of 3D organoid systems have been transplanted in vivo (Takebe et al., 2013; Watson et al., 2014), and in some cases they have been used to repair and rescue tissue damage (Yui et al., 2012), indicating that organoids have the potential to be good resources for cell therapy. Fred H Gage and colleagues proved that the intracerebral implantation of hiPSC-derived brain organoids into immunodeficient mice could develop into a functional vasculature system that supports the long-term survival of organoid grafts (Mansour et al., 2018). We further investigated whether the vascular systems in the vOrganoids could connect to host vascular systems to form a new functional circulation system when implanted in vivo. In the blood vessels that were located in the vOrganoid grafts regions, both human endothelial cells and mouse endothelial cells could be detected. Furthermore, a notable level of blood flow could be captured in the vOrganoid grafts by two-photon fluorescence imaging in mice. Therefore, our results illustrate that the vascular system in vOrganoids could recruit host endothelial cells to reconstruct functional vascular systems and to enable blood flow into the grafts after implantation. Vascularization is a vital factor for the survival of organoids that not only promotes the growth of cells in vitro but also plays a role in reconstructing blood vessels after transplantation in vivo. Hence, we speculate that our vascularized culture system may be widely applicable in future 3D organoid transplantations in vivo to improve the survival rates and functional reconstruction.

## Acknowledgements

This work was supported by the Strategic Priority Research Program of the Chinese Academy of Sciences (XDA16020601, XDB32010100), National Basic Research Program of China (2017YFA0103303, 2017YFA0102601), the National Natural Science Foundation of China (NSFC) (91732301, 31671072, 31771140, 81891001), the Grants of Beijing Brain Initiative of Beijing Municipal Science & Technology Commission (Z181100001518004).

## Author Contributions

Q.W. and X.W. conceived the project, designed the experiments and wrote the manuscript. Y. S., R. L., P. L. and L. G. performed the organoid culture. S. L. and R. C. performed electrophysiology analysis and calcium imaging. S. Z. and Y. S. carried out the single-cell RNA-seq experiment. Y. S., S. Z. and M. W. performed the RNA-seq data analysis. Y. S., Q.W., and A. F. performed immunefluorescent staining, imaging and analysis. Y. S. and J. L. performed transplantation, in vivo imaging and analysis. W. G. supervised angiogenesis analysis and evaluation. All authors edited and proofed the manuscript.

---

**Figure S1.**
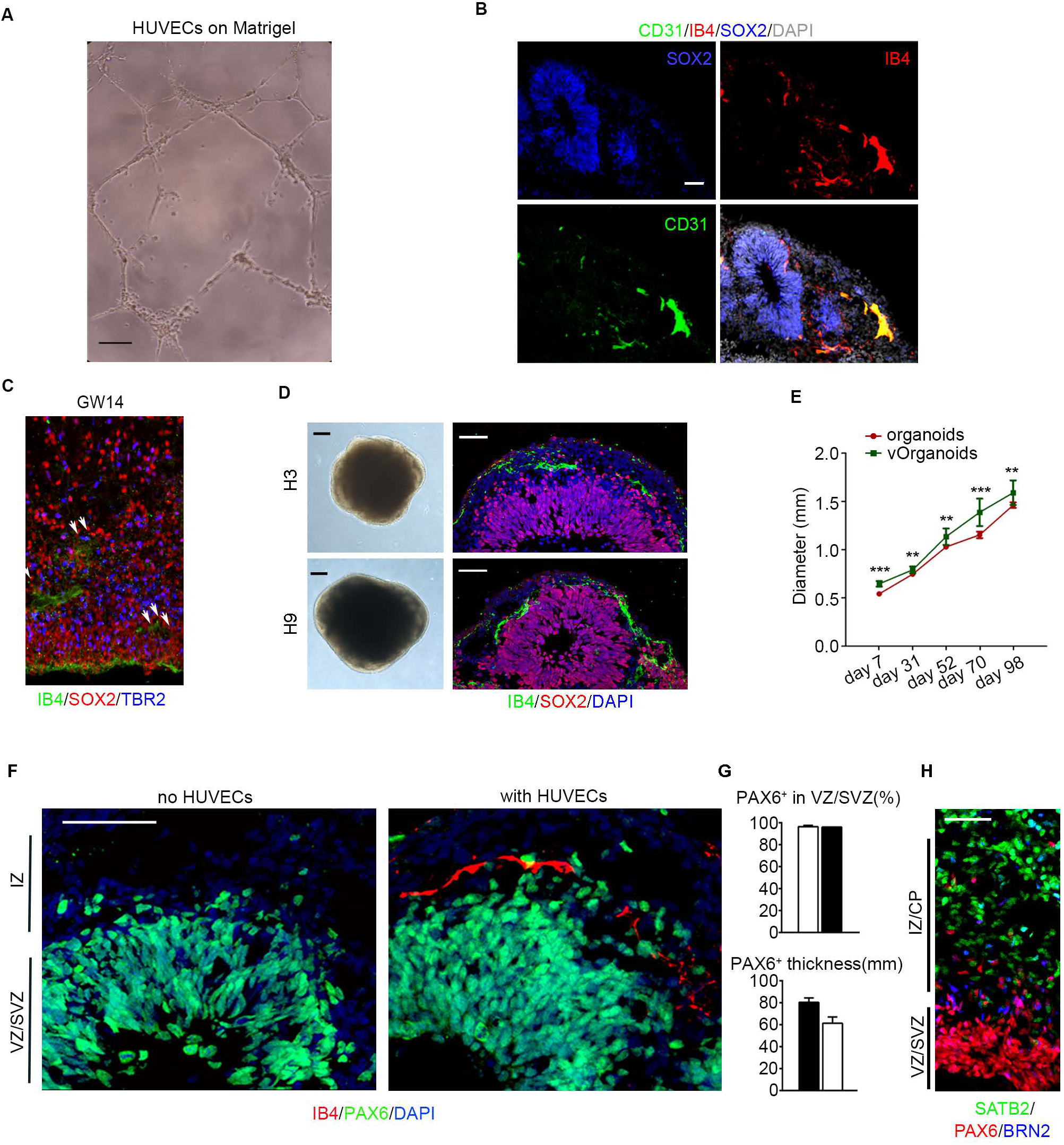
The vascular systems in vOrganoids promote the cell growth and reduce cell apoptosis. (A) Tube formation of HUVECs on matrigel. Scale bar, 200 μm. (B) Immunofluorescence staining for CD31, IB4 and SOX2 to illustrate that tube-like vascular systems were formed in vOrganoids at 45 days. Scale bar, 50 μm. (C) Staining for IB4 (green), SOX2 (Red), TBR2 (blue) in human cortical cryosections at GW14. Scale bar, 50 μm. (D) Bright field (BF) and immunohistochemical images of vOrganoids derived from hESC-3 and hESC-9 lines. Bright field images showed on the left and immunohistochemical images for vessel (IB4, green) and progenitor (SOX2, red) showed on the right. Scale bar, 50μm (left panels), 200 μm(right panels). (E) The diameters of organoids and vOrganoids generated from hESCs at day7, day30, day50, day70 and day100, respectively. (F) Distribution of progenitors (PAX6, green) in organoids with or without HUVECs (IB4, red). Scale bar, 50μm. (G) Quantification of percentages of PAX6^+^ cell within all cells (DAPI^+)^ in VZ/SVZ and of the thickness of PAX6^+^ region in organoids with or without HUVECs, respectively. n=3, 3 slices for Organoids with or without HUVECs. (H) Cryosections of vOrganoids were immunostained for progenitor and layer-specific cortical neuron markers at 65 days.

**Figure S2.**
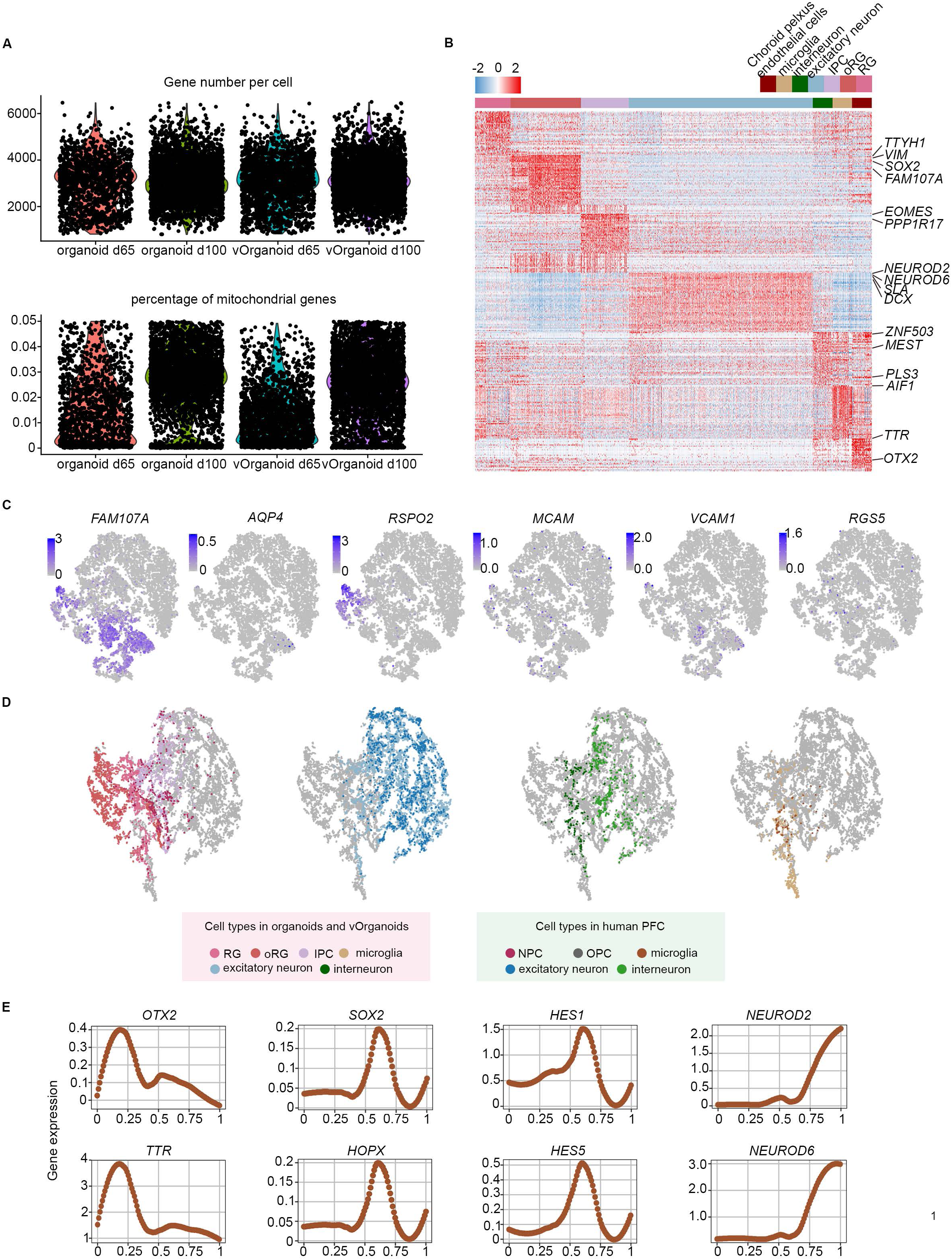
Single-cell RNA sequencing of organoids with and without HUVECs. (A) Violin plots demonstrated the gene number per cell and the percentage of mitochondrial genes of four organoids samples. (B) Heatmap showing the differential expressed genes for each cluster. Some canonical marker genes were labeled. (C) t-SNE visualizing of the expression of known marker genes. Cells are colored according to gene expression levels (blue, high; grey, low). (D) The correlations between organoids RG, oRG, IPC and PFC NPC; excitatory neurons derived from organoids and PFC ; interneurons derived from organoids and PFC; microglia derived from organoids and PFC were highlighted in tSNE, each dot was a single cell and colored by cell types. (E) Expression of canonical regulators, as a function of the position along the maturation trajectory. Curve reflects local averaging of single cell expression.

**Figure S3.**
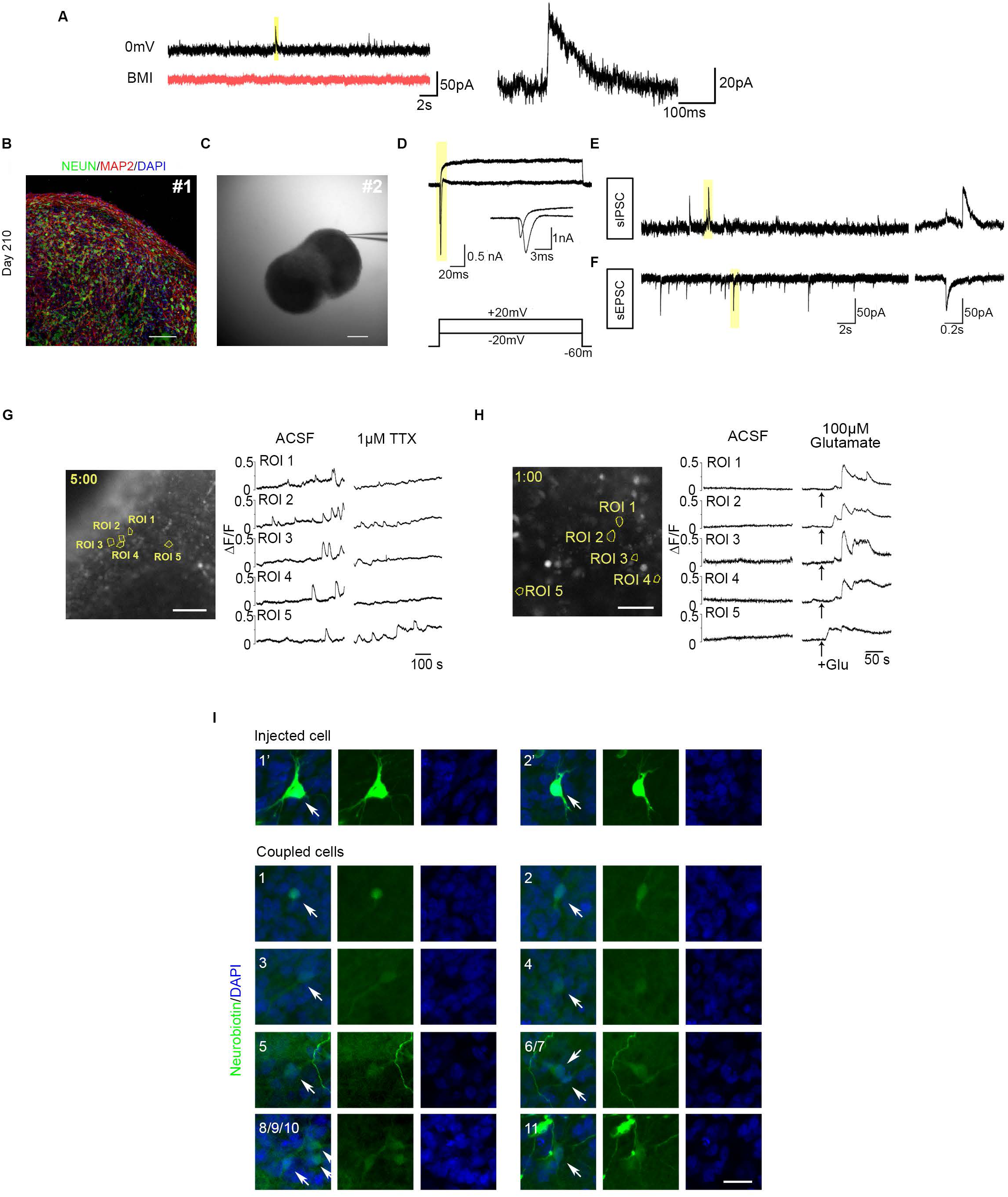
Functional circuits were built in vascularized organoids progressively. (A) Post-synaptic currents were sensitive to BMI. Representative IPSC event highlight by yellow bar displayed at the right panel. (B) Staining of classical neuron marker NeuN (green) and MAP2 (red) in the vOrganoid of day 210 (#1). Scale bar, 100 μm (C) An example of whole cell configuration in the vOrganoids of day 210 under DGC(#2). Scale bar, 200 μm (D) Representative traces of membrane currents elicited by depolarizing pulses of −20mV or +20mV from a neuron in the vOrganoid of day 210. (E-F) Spontaneous EPSCs and IPSCs recorded from the same cells in D. Yellow bars indicate representative events which are shown at right. (G-H) Intracellular spontaneous calcium fluctuations were measured before and after the application of 1 μM tetrodotoxin (TTX, G) or 100 μM glutamate (H) in the vOrganoids. Arrows mark the time of addition of glutamate. Scale bar: 100 μm in (G), 50 μm in (H). (I) Injected cells and the cells which coupled with them through gap-junctions were visualized by staining of neurobiotin.

**Figure S4.**
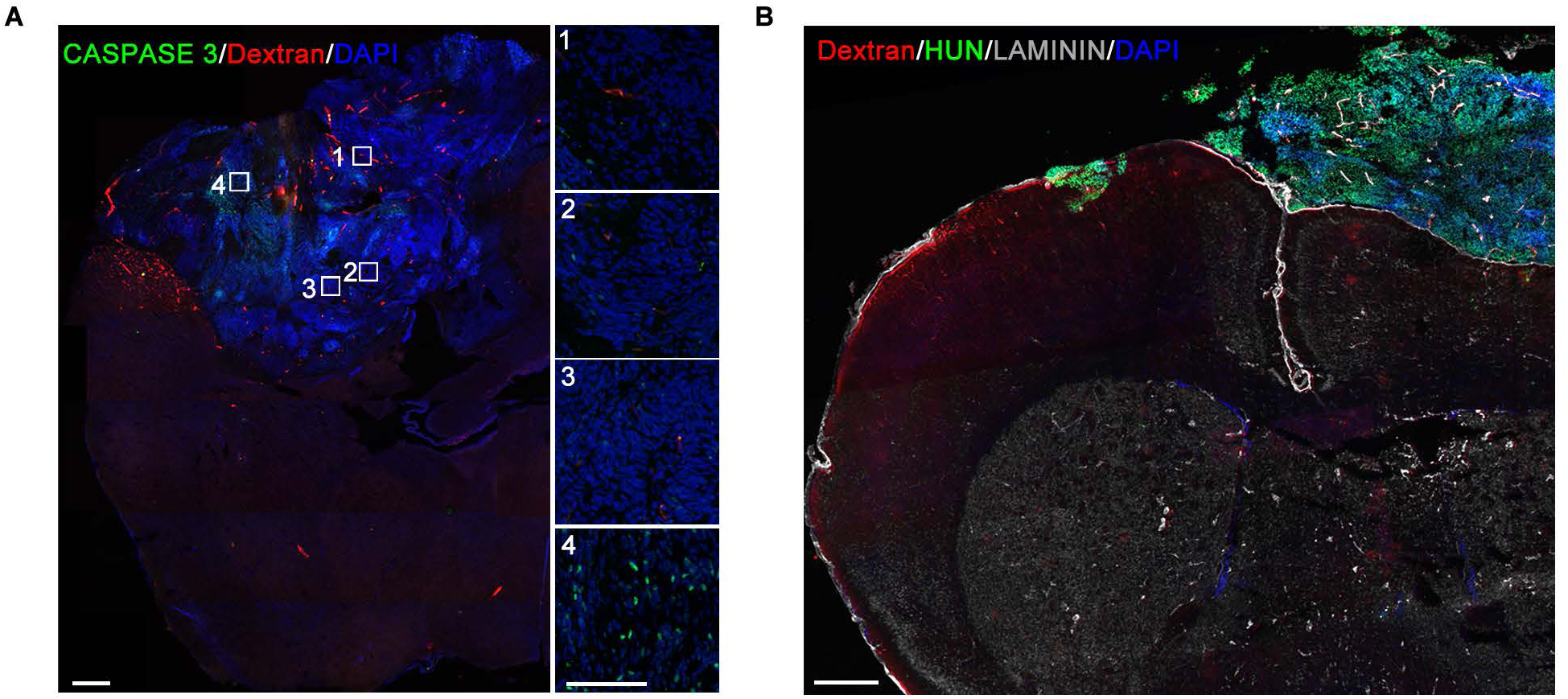
The vascular system formation in vOrganoids grafts. (A) Immunofluorescence staining for CASPASE 3 and Dextran in cryosections of host brain that contained the vOrganoid grafts. Scale bar, 500μm (left panels), 100 μm(right panels). (B) Immunofluorescence staining for laminin, HuN and Dextran in cryosections of host brain that contained the vOrganoid grafts to illustrate blood vessels formation in grafts at a global perspective.

**Movie S1** Movie shows the staining of LAMININ (red) and IB4 (green) displayed in 3D rotating around x,y,z-axis.

**Movie S2** Movie shows the 3D reconstruction of LAMININ-positive vascular structure rotating around y-axis.

**Movie S3** Movie shows the functional blood flow in vOrganoids grafts.

**Movie S4** Movie shows the 3D reconstruction of Dextran-positive blood vessels in vOrganoids grafts.

## STAR ★ Methods

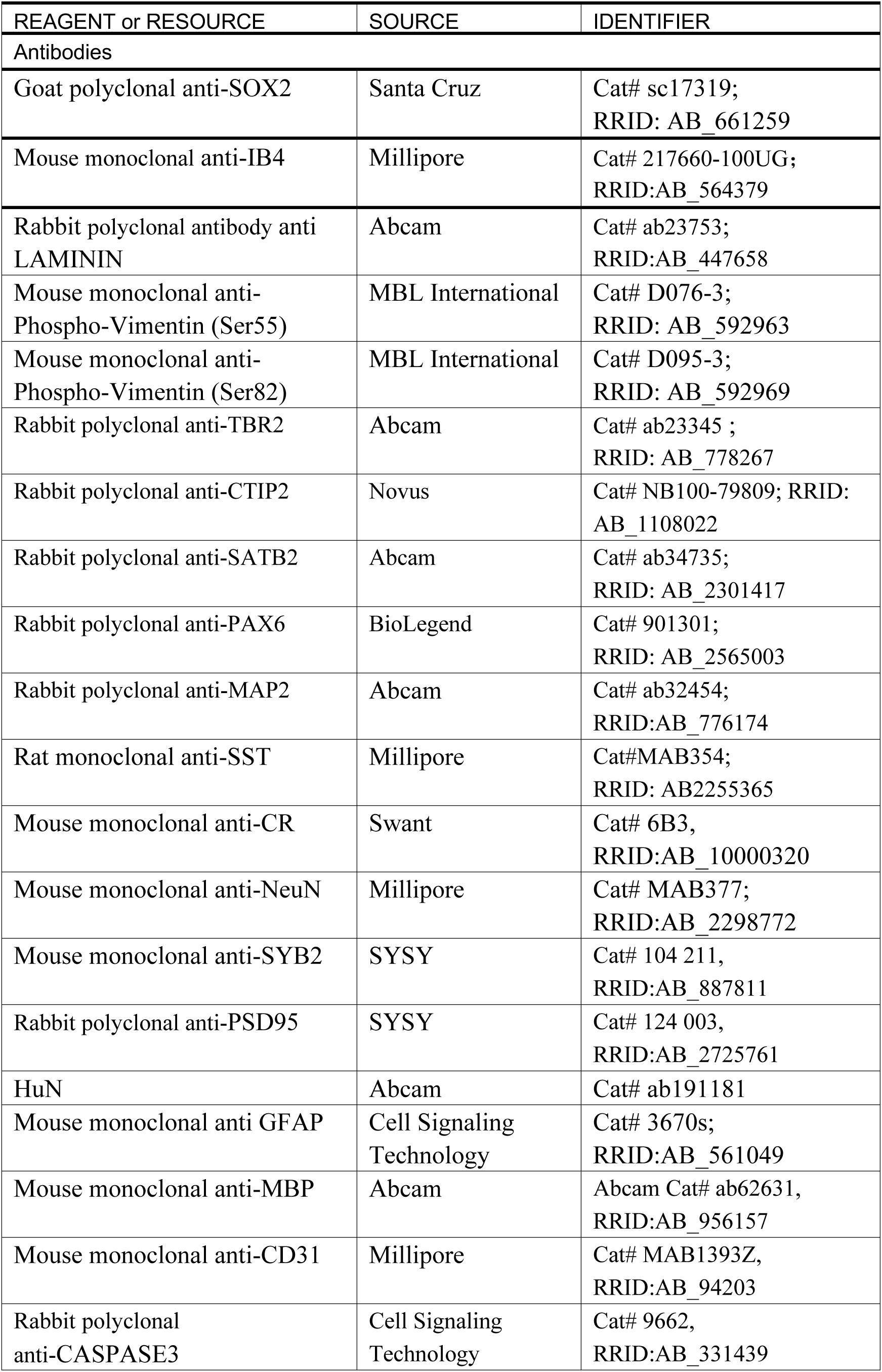
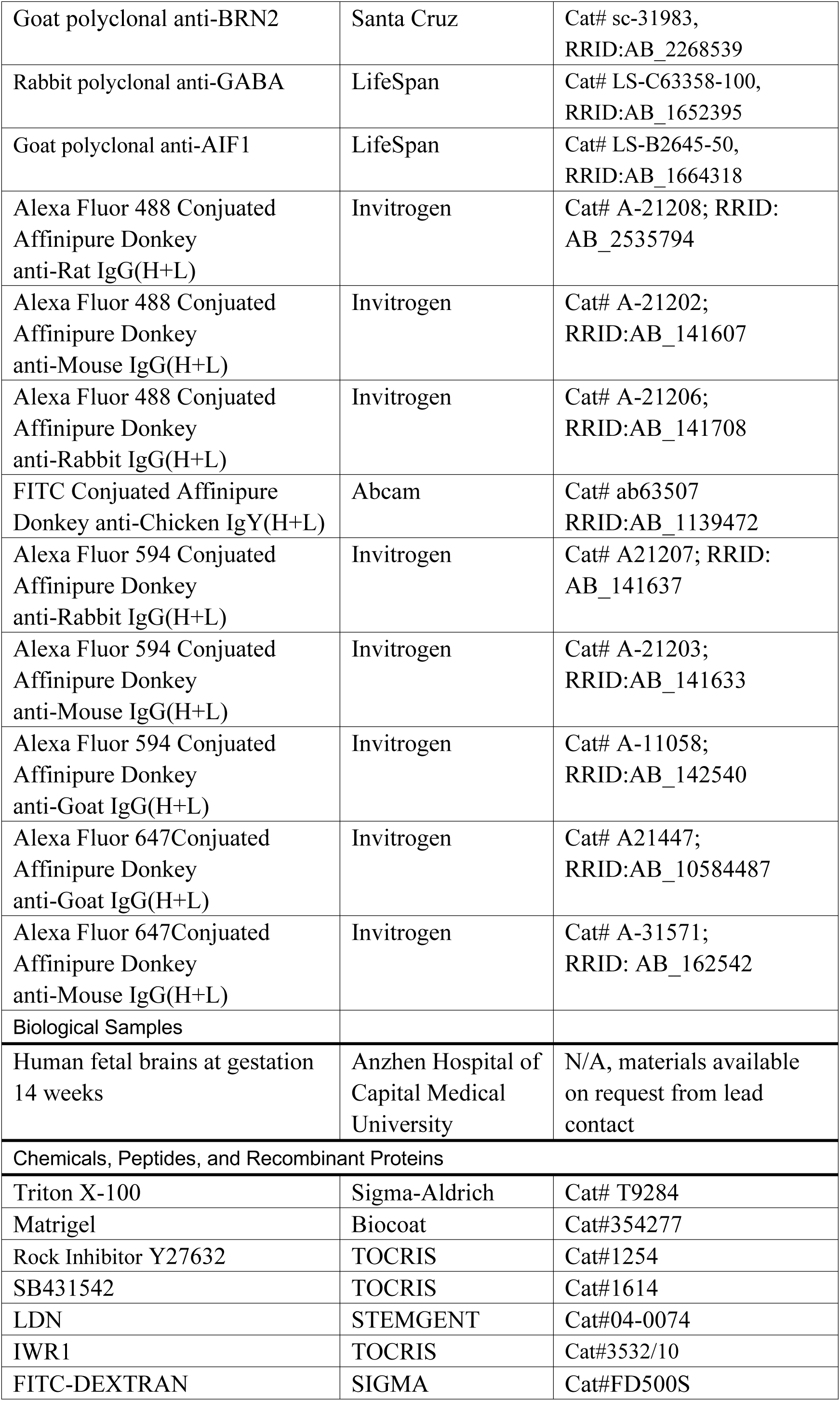
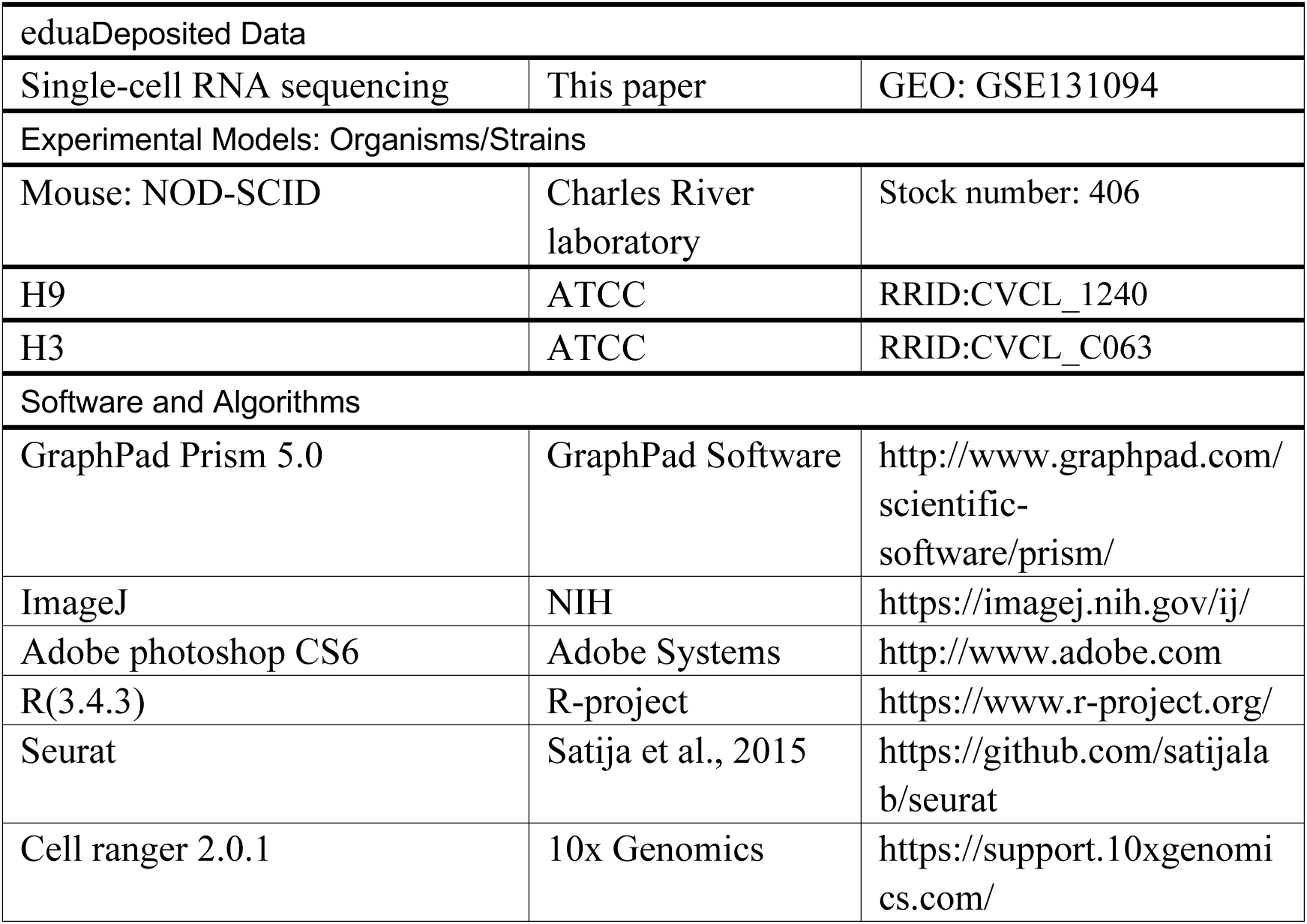
Key Resources Table.

| REAGENT or RESOURCE | SOURCE | IDENTIFIER |
| --- | --- | --- |
| Antibodies |  |  |
| Goat polyclonal anti-SOX2 | Santa Cruz | Cat# sc17319;<br>RRID: AB_661259 |
| Mouse monoclonal anti-IB4 | Millipore | Cat# 217660-100UG;<br>RRID:AB_564379 |
| Rabbit polyclonal antibody anti LAMININ | Abcam | Cat# ab23753;<br>RRID:AB_447658 |
| Mouse monoclonal anti-Phospho-Vimentin (Ser55) | MBL International | Cat# D076-3;<br>RRID: AB_592963 |
| Mouse monoclonal anti-Phospho-Vimentin (Ser82) | MBL International | Cat# D095-3;<br>RRID: AB_592969 |
| Rabbit polyclonal anti-TBR2 | Abcam | Cat# ab23345 ;<br>RRID: AB_778267 |
| Rabbit polyclonal anti-CTIP2 | Novus | Cat# NB100-79809; RRID: AB_1108022 |
| Rabbit polyclonal anti-SATB2 | Abcam | Cat# ab34735;<br>RRID: AB_2301417 |
| Rabbit polyclonal anti-PAX6 | BioLegend | Cat# 901301;<br>RRID: AB_2565003 |
| Rabbit polyclonal anti-MAP2 | Abcam | Cat# ab32454;<br>RRID:AB_776174 |
| Rat monoclonal anti-SST | Millipore | Cat#MAB354;<br>RRID: AB2255365 |
| Mouse monoclonal anti-CR | Swant | Cat# 6B3,<br>RRID:AB_10000320 |
| Mouse monoclonal anti-NeuN | Millipore | Cat# MAB377;<br>RRID:AB_2298772 |
| Mouse monoclonal anti-SYB2 | SYSY | Cat# 104 211,<br>RRID:AB_887811 |
| Rabbit polyclonal anti-PSD95 | SYSY | Cat# 124 003,<br>RRID:AB_2725761 |
| HuN | Abcam | Cat# ab191181 |
| Mouse monoclonal anti GFAP | Cell Signaling Technology | Cat# 3670s;<br>RRID:AB_561049 |
| Mouse monoclonal anti-MBP | Abcam | Abcam Cat# ab62631,<br>RRID:AB_956157 |
| Mouse monoclonal anti-CD31 | Millipore | Cat# MAB1393Z,<br>RRID:AB_94203 |
| Rabbit polyclonal anti-CASPASE3 | Cell Signaling Technology | Cat# 9662,<br>RRID:AB_331439 |
| Goat polyclonal anti-BRN2 | Santa Cruz | Cat# sc-31983,<br>RRID:AB_2268539 |
| Rabbit polyclonal anti-GABA | LifeSpan | Cat# LS-C63358-100,<br>RRID:AB_1652395 |
| Goat polyclonal anti-AIF1 | LifeSpan | Cat# LS-B2645-50,<br>RRID:AB_1664318 |
| Alexa Fluor 488 Conjugated<br>Affinipure Donkey<br>anti-Rat IgG(H+L) | Invitrogen | Cat# A-21208; RRID:<br>AB_2535794 |
| Alexa Fluor 488 Conjugated<br>Affinipure Donkey<br>anti-Mouse IgG(H+L) | Invitrogen | Cat# A-21202;<br>RRID:AB_141607 |
| Alexa Fluor 488 Conjugated<br>Affinipure Donkey<br>anti-Rabbit IgG(H+L) | Invitrogen | Cat# A-21206;<br>RRID:AB_141708 |
| FITC Conjugated Affinipure<br>Donkey anti-Chicken IgY(H+L) | Abcam | Cat# ab63507<br>RRID:AB_1139472 |
| Alexa Fluor 594 Conjugated<br>Affinipure Donkey<br>anti-Rabbit IgG(H+L) | Invitrogen | Cat# A21207; RRID:<br>AB_141637 |
| Alexa Fluor 594 Conjugated<br>Affinipure Donkey<br>anti-Mouse IgG(H+L) | Invitrogen | Cat# A-21203;<br>RRID:AB_141633 |
| Alexa Fluor 594 Conjugated<br>Affinipure Donkey<br>anti-Goat IgG(H+L) | Invitrogen | Cat# A-11058;<br>RRID:AB_142540 |
| Alexa Fluor 647 Conjugated<br>Affinipure Donkey<br>anti-Goat IgG(H+L) | Invitrogen | Cat# A21447;<br>RRID:AB_10584487 |
| Alexa Fluor 647 Conjugated<br>Affinipure Donkey<br>anti-Mouse IgG(H+L) | Invitrogen | Cat# A-31571;<br>RRID: AB_162542 |
| Biological Samples |  |  |
| Human fetal brains at gestation<br>14 weeks | Anzhen Hospital of<br>Capital Medical<br>University | N/A, materials available<br>on request from lead<br>contact |
| Chemicals, Peptides, and Recombinant Proteins |  |  |
| Triton X-100 | Sigma-Aldrich | Cat# T9284 |
| Matrigel | Biocoat | Cat#354277 |
| Rock Inhibitor Y27632 | TOCRIS | Cat#1254 |
| SB431542 | TOCRIS | Cat#1614 |
| LDN | STEMGENT | Cat#04-0074 |
| IWR1 | TOCRIS | Cat#3532/10 |
| FITC-DEXTRAN | SIGMA | Cat#FD500S |
| eduaDeposited Data |  |  |
| Single-cell RNA sequencing | This paper | GEO: GSE131094 |
| Experimental Models: Organisms/Strains |  |  |
| Mouse: NOD-SCID | Charles River laboratory | Stock number: 406 |
| H9 | ATCC | RRID:CVCL_1240 |
| H3 | ATCC | RRID:CVCL_C063 |
| Software and Algorithms |  |  |
| GraphPad Prism 5.0 | GraphPad Software | http://www.graphpad.com/scientific-software/prism/ |
| ImageJ | NIH | https://imagej.nih.gov/ij/ |
| Adobe photoshop CS6 | Adobe Systems | http://www.adobe.com |
| R(3.4.3) | R-project | https://www.r-project.org/ |
| Seurat | Satija et al., 2015 | https://github.com/satijalab/seurat |
| Cell ranger 2.0.1 | 10x Genomics | https://support.10xgenomics.com/ |

### Contact for Reagent and Resource Sharing

Further information and requests for resources and reagents should be directed to and will be fulfilled by the Lead Contact, Xiaoqun Wang.

### Experimental Model and Subject Details

#### Mice

NOD-SCID- immunodeficient mice were purchased from Charles River Laboratories in China (Vital river, Beijing, China) and used for experiments of organoids implantations. The animal housing conditions and all experimental procedures in this study were in compliance with the guidelines of the Institutional Animal Care and Use Committee of the Institute of Biophysics, Chinese Academy of Sciences. All mice were housed in SPF environments with a 12 hr light-dark schedule and had free access to food and water. All the subjects were not involved in any previous procedures.

#### Cell lines

Cell lines including hESCs H3 and H9 were purchased from ATCC and used for 3D organoids culture. Research protocol was approved by the institutional review board (ethics committee) of the Institute of Biophysics, Chinese Academy of Sciences.

#### Tissue sample collection

The human brain tissue collection and research protocols were approved by the Reproductive Study Ethics Committee of Beijing Anzhen Hospital and the Institute of Biophysics. The informed consent was designed as recommended by the ISSCR guidelines for fetal tissue donation and the fetal tissue samples were collected after the donor patients signing informed consent document which is in strict observance of the legal and institutional ethical regulations from elective pregnancy termination specimens at Beijing Anzhen Hospital, Capital Medical University. All the protocols are in compliance with the “Interim Measures for the Administration of Human Genetic Resources”, administered by the Chinese Ministry of Health. Gestational age was measured in weeks from the first day of the woman’s last menstrual cycle to the sample collecting date. Fetal brain were collected in ice cold artificial cerebrospinal fluid containing 125.0 mM NaCl, 26.0 mM NaHCO3, 2.5 mM KCl, 2.0mM CaCl2, 1.0 mM MgCl2, 1.25 mM NaH2PO4 at a pH of 7.4 when oxygenated (95% O2 and 5% CO2).

### Method Details

#### 3D vOrganoids culture and differentiation procedure

hESCs H9 and H3 lines were maintained on Matrigel-coated 6-well plates (Corning) and were cultured with Essential 8™ Medium (Gibco). On day 0, hESCs colonies were pretreated for one hour with 20 µM Y27632 (Tocris Bioscience), and were dissociated into single cell by Accutase (STEMCELL Technologies). HUVECs were cultured in endothelial Cell Growth Medium, and dissociated into single cell by TrpLE. The mixture of approximately 3×10^6^ dissociated hESCs and 3×10^5^ HUVECs were resuspended in KSR medium, and plated into 96-well U shaped polystyrene plates (Thermo Fisher).The size of EBs is determined by the cell numbers that seeding into each well of the plate. The KSR medium was prepared as follows: DMEM/F12 (Gibco) was supplemented with 20% Knockout Serum Replacement (KSR, Gibco), 2 mM Glutamax (Gibco), 0.1mM nonessential amino acids (NEAA, Gibco), 0.1 mM beta-mercaptoethanol (Gibco), 3 μM endo-IWR1 (Tocris Bioscience), 0.1 μM LDN-193189 (STEMGENT) and 10 μM SB431542 (Tocris Bioscience). On day 18, the self-organized floating EBs were transferred to low-cell-adhesion 6-well plates (Corning) and further cultured in the neural induction medium containing DMEM/F12, 1:100 N2 supplement (Gibco), 2 mM Glutamax (Gibco), 0.1 mM nonessential amino acids (NEAA, Gibco), 55 μM beta-mercaptoethanol (Gibco), 1 μg/ml heparin. After day35, the free-floating aggregates were transferred to the neurobasal-type differentiation medium supplemented with 1:50 B27 (Gibco), 2 mM Glutamax and 0.1 mM NEAA, 0. 55 μM beta-mercaptoethanol, 5 μg/ml heparin, 1% matrigel, 10 ng/ml BDNF, 10 ng/ml GDNF, 1 μM cAMP (Sigma).

#### Immunofluorescence

Organoids and tissues were fixed by 4% paraformaldehyde in PBS at 4°C for 2h, and then dehydrated in 30% sucrose in PBS. The fixed and dehydrated organoids and tissues were embedded and frozen at -80°C in O.C.T. compound, sectioned with Leica CM3050S. Cryosections were subjected to antigen retrieval, pretreated (0.3% Triton X-100 in PBS) and incubated for a blocking solution (10% normal donkey serum, 0.1% Triton X-100, and 0.2% gelatin in PBS), followed by incubation with the primary antibodies overnight at 4°C. Immunofluorescence images were acquired with Olympus laser confocal microscope and analyzed with FV10-ASW viewer (Olympus), ImageJ (NIH) and Photoshop (Adobe).

#### Electrophysiology

Cultured vOrganoids were embedded in 3% low-melting agarose in ACSF (in mM: 126 NaCl, 3 KCl, 26 NaHCO_3_, 1.2 NaH_2_PO_4_, 10 D-glucose, 2.4 CaCl_2_ and 1.3 MgCl_2_) and sectioned at 200 μm in oxygenated (95% O2 and 5% CO2) ice-cold ACSF with a vibratome (VT1200s, Leica). The slices were then cultured in a 24-well plate filled with 250 μl/well of neural differentiation medium in an incubator (5% CO_2_, 37°C). After a recovery period of at least 24 hours, an individual slice was transferred to a recording chamber and continuously superfused with oxygenated ACSF at a rate of 3-5 mL per minute at 30±1°C. Whole-cell patch clamp recording was performed on cells of vOrganoid slices. Patch pipettes had a 5-7 MΩ resistance when filled with intracellular solution (in mM: 130 potassium gluconate, 16 KCl, 2 MgCl_2_, 10 HEPES, 0.2 EGTA, 4 Na_2_-ATP, 0.4 Na_3_-GTP, 0.1% Lucifer Yellow and 0.5% neurobiotin, pH = 7.25, adjusted with KOH). Evoked action potentials were recorded in current-clamp mode using a series of injected currents from −50 pA to 300 pA in increments of 50 pA. Whole-cell currents were recorded in voltage-clamp mode with a basal holding potential of −60 mV followed by stimulating pulses from −80 mV to 40 mV with a step size of 10 mV. The membrane potential was held at −70/0 mV when spontaneous EPSCs/IPSCs were recorded. Dual-patch recording was performed in current-clamp mode. A pair of pulses (1nA, 2ms duration, 50ms interval) were injected into each cell separately. The cells were monitored with a 40x Olympus water-immersion objective lens, a microscope (Olympus, BX51 WI) configured for DGC and a camera (Andor iXon3). Stimulus delivery and data acquisition were conducted with a multiclamp 700B amplifier and a Digidata 1440A (Molecular Devices), which were controlled by Clampex 10. The slices were fixed after patch clamp recording. Staining with fluorescein streptavidin (Vector SA-5001, 1:500) or Texas Red streptavidin (Vector SA-5006, 1:500) was performed to visualize the morphology of cells.

#### Calcium imaging

3 μl of dye solution that contained 50 μg Fluo-4 AM (Life Technologies), 50 μL DMSO and 200 μg Pluronic F-127 (Sigma) was applied to the surface of each individual slice. After incubation for 30 min at 37°C, the slice was transferred to a recording chamber and continuously superfused with oxygenated ACSF (in mM: 126 NaCl, 3 KCl, 26 NaHCO3, 1.2 NaH2PO4, 10 D-glucose, 2.4 CaCl2 and 1.3 MgCl2) at a rate of 3-5 mL per minute at 30±1°C. The slices were washed 30 minutes before imaging. Calcium imaging was acquired at 5 Hz using a camera (Andor iXon3) with a FITC filter set (Ex: 475/35 nm, Em: 530/43 nm) on a BX51WI microscope (Olympus). Data analysis was performed with ImageJ. The ROIs were selected manually, and the mean fluorescence (F) was calculated for each frame. The fluorescence changes over time was calculated as follows: ΔF=(F-Fbasal)/Fbackground, in which Fbasal was the lowest mean fluorescence value during imaging, and Fbackground was the average mean fluorescence across all frames. Since 10 minutes before imaging, TTX (1 μM), CNQX (20 μM), APV (100 μM) and BMI (20 μM) were added by bath application. The slices were rinsed for 30 minutes with ACSF after drug treatments. Different from other drugs, glutamate (100 μM) was added by bath application during imaging.

#### Implantation of cerebral organoid into mice S1 cortex

Before implantation, cerebral organoids had been cultured for 60 days. And strict screening was performed by brightfield microscopy to select the organoids with appropriate size and displayed without massive cyst formation. Immune-deficient NOD-SCID mice, aged 8 weeks were used in our studies. Mice were anesthetized by intraperitoneal injection of avertin. The heads of animals were fixed in a stereotactic frame. The fur above the skull was removed and skin was cut. A∼3-mm diameter craniotomy was performed by polishing the skull; the underlying dura mater was removed and a cavity was made by aspiration with a blunt-tip needle attached to a vacuum line. The aspirative lesion was made unilaterally in the region of the S1cortex.

Sterile ACSF was used to irrigate the lesion and keep it free of blood throughout the surgery, and a piece of Gelfoam (Pfizer) was used to slow the bleeding and absorb the excess blood. Molten 3% low-melting agarose was dropped in the implantation regions to immobilize the organoid grafts as agarose congealed quickly. Adhesive glue was used to seal the border of implanted organoid grafts. The wound was closed with sutures. Following completion of the surgery, penicillin streptomycin combination was administrated for inflammation and analgesic relief. The mice were then returned into home cages to recover.

#### Two- photon imaging

For imaging of blood flow in organoid grafts, mice were tail intravenously injected FITC-Dextran. The mouse was fixed on the recording setup, with an isoflurane-oxygen mixture of 0.5-1% (v/v). The in vivo imaging of blood flow was done with a 2-photon laser scanning microscope. The recording chamber was perfused with normal artificial cerebral spinal fluid (ACSF) containing 126 mM NaCl, 3 mM KCl, 1.2 mM NaH_2_PO_4_, 2.4 mM CaCl_2_, 1.3 mM MgCl_2_, 26 mM NaHCO_3_, and 10 mM D-glucose (pH 7.4 when bubbled with 95% oxygen and 5% CO_2_). The temperature of the mouse was kept at ∼37°C throughout the experiment.

#### Single-Cell Dissociation, and library construction

The organoids with and without HUVECs at 65 days and 100 days, respectively, were cut into small pieces and dissociated into single-cell suspensions by using a papain-based dissociation protocol (hibernate E medium with 1 mg/ml papain (Sigma) at 37°C on a thermo cycler with 500g for 15-20 min). Single cells were suspended in 0.04% BSA/PBS at the proper concentration to generate cDNA libraries with Single Cell 3’ Reagent Kits, according to the manufacturer’s protocol. Briefly, after the cDNA amplification, enzymatic fragmentation and size selection were performed to optimize the cDNA size. P5, P7, an index sample, and R2 (read 2 primer sequence) were added to each selected cDNA during end repair and adaptor ligation. P5 and P7 primers were used in Illumina bridge amplification of the cDNA (http://10xgenomics.com). Finally, the library was processed on the Illumina platform for sequencing with 150bp pair-end reads.

#### Data processing of single-cell RNA-seq

Cell ranger 2.0.1 was used to perform quality control and read counting of *Ensemble* genes with default parameter (v2.0.1) by mapping to the GRCh38 human genome. We excluded poor quality cells by the Seurat package (v2.1.0) in Bioconductor after the gene-cell data matrix was generated by cell ranger. Only cells that expressed more than 800 genes were considered, and only genes expressed in at least 200 single cells were included for further analysis. Besides, the cells of percentage of mitochondrial genes above 0.05 were also omitted in subsequent analysis.

#### Identification of cell types in organoids by dimensional reduction

Seurat (v2.1.0) was used to perform linear dimensional reduction. 3805 highly variable genes with an average expression between 0.0125 and 3 and dispersion greater than 0.5 were selected as input for principal component analysis (PCA). The significant PCs were used for 2-dimension tSNE to cluster cells by the FindClusters function with Resolution 6.5. Clusters were identified by the specific expression of known markers and GO analysis.

#### Identification of differentially expressed genes among clusters

Genes differentially expressed in each cluster were identified using the FindAllMarkers function (thresh.use =0.25) in the Seurat R package. The genes with average expression difference > 0.25 natural log with p < 0.05 were selected as marker genes. Gene Ontology terms were obtained using DAVID 6.7.

#### Mapping organoids cell types to PFC cell types

Organoid and PFC datasets were mapped based common sources of variation. We first identified top 1000 highly variable genes (HVGs) in each dataset using function FindVariableGenes with default parameters. We then created a union of common HVGs, which was used as input to the canonical correlation analysis (RunCCA function). We selected the top 10 canonical correlation vectors which had a minimum bicor of 0.2 to align these two datasets by applying function AlignSubspace.

To further investigate the correlation between organoid cell types and PFC cell types, we first computed the coordinate of centroid of each cluster, then the distance and correlation between each two clusters. The cluster-cluster network was constructed by function igraph (http://igraph.org)(Csardi and Nepusz) and visualized using Cytoscpae (Shannon et al., 2003), intra-cluster edges and edges with correlation coefficient smaller than 0.3 were removed. Edge width and color correspond to the inter-cluster correlation.

#### Identification of maturation trajectory

To order cells along maturation trajectory, we first performed dimensionality reduction using diffusion maps, then fit a principle curve through the data using R Package princurve (http://CRAN.R-project.org/package=princurve) (Weingessel and Hastie, 2007). The arc-length from the beginning of the curve to the point where the cell projects onto the curve was defined as the maturation of each cell. To assign the direction of the resulting curve, we correlated the expression of gene OTX2 with maturation. Cells with negative correlation were ordered at the beginning of the curve. OTX2 is a known choroid plexus related gene and thus should has high expression level at the beginning of the curve.

#### Quantification and Statistical Analysis

All data were represented as the mean ± SEM. The quantification graphs were made using GraphPad Prism software. Sample size (n) for each analysis can be found in the figure legends.

#### Data Resources

The accession number for the RNA sequencing data reported in this paper is GEO: GSE131094.

